# AHR agonist ITE boosted PD1 antibody’s effects by inhibiting myeloid-derived cells suppressive cells in an orthotopic mouse glioma model

**DOI:** 10.1101/2022.03.14.484219

**Authors:** Lijiao Zhao, Yunlong Ma, Qiuting Shu, Hui Sun, Jing Lu, Pei Gong, Fanhua Meng, Fang Wan

## Abstract

Glioblastoma is a “cold“ tumor lacking T cell infiltration and tryptophan metabolites such as kynurenine function as an immune suppressor by binding to aryl hydrocarbon receptor(AHR). Hence AHR antagonists have been developed and some shown to activate the immune response in IDO over-expressing cancer models. Paradoxically, AHR has been reported to block glioma cell invasion like a tumor suppressor, and how to target AHR in cancer remains an open question. We previously discovered that an AHR agonist ITE can effectively inhibit glioma invasion. Here we report that ITE combined with PD1 antibody significantly increased the infiltration of CD8+ T cells while reducing that of the myeloid-derived suppressive cells (MDSCs), extending mouse survival. To identify factors that possibly mediated ITE’s effects, we analyzed RNA-seq data and discovered that ITE significantly down-regulated IL11, a known MDSC regulator. In contrast, kynurenine upregulated IL11, further supporting IL11 as an AHR target. Moreover, ITE inhibited IL11’s induction of MDSC from mouse PBMC in vitro, supporting that IL11 might mediate ITE’s MDSC regulation effects. The discovery of ITE’s immune-activating effects highlighted the complexity of AHR’s signaling in cancers. The unexpected increase in STAT3, downstream of IL11 provided clues for combination therapy development.

## 1 Introduction

Immunotherapy for GBM has been challenging, because the majority of GBM cancers are “cold” tumors with less cytotoxic immune cell infiltration, although the total immune cell infiltration is more in glioma compared to the normal brain^[1]^Systematic immune function was inhibited in both GBM patient and the mouse orthotopic models, with impaired T cells function and more MDSC^[2–4]^. Moreover, T cells were sequestered into the bone marrow from periphery blood in GBM patients^[5]^.

GBM cells express proteins on cell surfaces that directly inhibit cytotoxic immune cells, such as HLA-G, HLA-E for NK cells inhibition, and PDL1 for inhibition of CD8+ T cells ^[6]^. GBM cells also release immune-suppressing proteins and molecules into the tumor microenvironment, such as KYN and TGF beta^[7, 8]^, exosomes containing PD1 protein, etc. Kyurenine is a tryptophan metabolite generated by IDO, which was overexpressed in glioblastoma specimens and significantly associated with a poor prognosis^[9]^. Furthermore, IFN-γ can induce IDO expression^[10]^, which was responsible for mediating the adaptive resistance of tumors to PD-1/PD-L1 or CTLA-4 checkpoint blockades^[11]^.

To turn the tide of the highly suppressive GBM microenvironment requires activation of both innate and adaptive immunity, and mere checkpoint blockade is not enough. Anti PD1 antibody nivolumab in GBM patients was in phase III clinical trial, but preliminary results showed that endpoint was not met: median overall survival in the nivolumab treatment group is similar to the bevacizumab treated control group (NCT02017717)^[12]^.

One major PD1 resistance pathway is the IDO/TDO-Kyurenine-AHR pathway, and the patients’ KYN/Trp ratio is prognostic for GBM immunotherapy outcome^[13]^. However, a combination of PD1 blockade with IDO/TDO inhibitor failed to meet the expectation in other cancer types in clinical trials(NCT03707457). Adding radiation to IDO1 inhibitor and PD1 blockade increased survival in the orthotopic murine glioma model^[14]^, supporting that IDO inhibition alone is not sufficient for overcoming PD1 resistance in glioma. Recently it’s found that IDO/TDO is not the only immune-suppressive pathway via AHR, as another IDO/TDO independent tryptophan metabolite, indole-3-acetate, binds to AHR and suppress immunity too^[15]^.

Directly targeting AHR is probably a better strategy for overcoming PD1 antibody resistance, and several AHR antagonists are in development. However, AHR acted as a tumor suppressor in glioma by inhibiting glioma cell invasion, and we previously discovered that a high-affinity AHR agonist, ITE, can block glioma invasion^[16]^. Moreover, ITE bound AHR inhibited transcription of POU5F1 in glioma stem cells, and inhibited the tumor growth in the U87MG xenograft glioma model^[17]^. AHR has also been recently identified as a suppressor of lung metastasis^[18]^. We hypothesized that ITE can compete with endogenous AHR ligand kynurenine and indole-3-acetate, and overcome PD1 antibody resistance. Here we reported that the combination of ITE and PD1 antibody increased CD8+ T cells infiltration in an orthotopic glioma model, probably via MDSC inhibition, and extended the mouse survival. Inhibition of glioma cells’ IL11 expression was identified to be a possible mediator of ITE’s effects.

## 2 Materials and Methods

### 2.1 Cell lines, animals, and reagents

Mouse glioma cell lines GL261 were kindly provided by the Third Military Medical University. GL261 cells were cultured using DMEM/F12 (Gibco, 8121048) supplemented with 10% FBS (Tianhang, 22011-8612), 6-to 8-week-old C57BL/6J wild-type female mice were purchased from Beijing Vital River Laboratory Animal Technology (Co. Ltd). All animal experiments were performed per protocols approved by the animal care guidelines of the Inner Mongolia Agricultural University. PD1 antibody (BioXCell, BE0273) and other antibodies were listed in the supplemental table 1.

### 2.2 Mouse orthotopic glioma model

Mice were anesthetized with an intraperitoneal injection of 0.3% Pentobarbital sodium (30mg/kg). For the stereotactic intracranial injection, the surgical site was shaved and prepared with iodine. A midline incision was made to expose the bregma point, 1×105 GL261 cells in a volume of 5ul medium were stereotactically injected into the left striatum defined by the following coordinates: 2 mm posterior to the coronal suture, 2 mm lateral to the sagittal suture, and 3mm deep to the cortical surface, using a microinjection pump (KD scientific, 78-1311Q) at a speed of 0.5ul/min. The needle was removed slowly and the skin was sutured with nylon thread.

### 2.3 Drug treatment

The mice were randomly divided into four groups (n = 8 mice/group): control, ITE group, anti-PD1 monoclonal antibody treatment (PD1 group), and ITE plus anti-PD1 monoclonal antibody (ITE+PD1 group). The anti-PD1 monoclonal antibody was given by intraperitoneal injection twice, with each injection at 100ug, separated by three days. ITE was dissolved in 5% DMSO, i.p.at 100 mg/kg, given every other day. The ITE+PD1 group was given 100mg/kg ITE two times, separated by one day, then given 100ug PD1 once. The control group was given the same volume of 5% DMSO solution at the same schedule as the ITE group.

### 2.4 Animal imaging

IR783 dye was utilized for in vivo imaging of the glioma[19]. Mice bearing orthotopic tumors were injected intraperitoneally with IR-783 at a dose of 50 nmol/mouse 24hr before imaging. Imaging was done on IVIS Lumina XR Imaging System (PerkinElmer, Waltham, MA) equipped with fluorescent filter sets (excitation/emission, 783/840 nm), with automatic subtraction of background fluorescence from the total dye uptake in tumors during the image acquisition, as described previously[14, 22], on 11, 14, 17, 20, 23 and 26 days post tumor implantation. The mice were euthanized when the posterior tumor edge reaches the tragus of the ears, judged by in vivo imaging, to ensure that tumor size is below 20mm for animal welfare, and survival was analyzed by Kaplan-Meier analysis using IBM SPSS 20.0 software. For tumor growth analysis, all animals were imaged from each control or treatment arm, and tumor volume data were analyzed with the TumorGrowth online tool.

### 2.5 Preparation of single-cell suspension

For subsequent analysis by flow cytometry, tumors were cut into pieces and digested with Collagenase IV (200U/ml) and DNase I (100mg/ml) for 30min at 37°C. Tissue was passed through a 70mm cell strainer (Falcon) and washed with PBS buffer before proceeding with antibody-mediated staining.

### 2.6 Flow Cytometry

Flow cytometry experiments were carried out on the Beckman CytoFLEX S flow cytometer, and data were analyzed using Beckman CytoFLEX S software. For all experiments, cell debris was excluded by forwarding versus side scatter plot, and doublets were excluded by scattering height vs forward scatter area plot. Nonviable cells were excluded by Live/Dead Aqua (Invitrogen) staining. Fluorescent compensation experiments were performed by staining each antigen separately to generate the compensation matrix.

For the multi-color immune cell analysis, single-cell suspension per brain tumor sample was firstly stained with Aqua Live/Dead viability dye (Biolegend) according to the manufacturer’s instructions. Cells were then incubated in blocking solution TruStain FcX™ (anti-mouse CD16/32) Antibody (Bio legend 101320) in PBS, and then stained with a standard panel of immunophenotyping antibodies in staining buffer (See Table for a list of antibodies, fluorochromes, manufacturers, and concentrations) for 30 minutes at 4°C. After staining, cells were washed with PBS, then fixed in 1:3 fixation/permeabilization concentrate: diluent mixture (BD) for 15 minutes, centrifuged, washed, then stained for FoxP3, IL17 in permeabilization buffer for 30 minutes at 4°C. Cells were then centrifugated at 450g, and the pellet was resuspended in staining buffer before loading into the flow cytometer.

For the MDSC induction analysis, normal mouse peripheral blood single cells were collected and stained with each antibody separately, then analyzed with the flow cytometer.

### 2.7 Immunohistochemistry (IHC)

The brains were extracted from the mice and fixed with 4% paraformaldehyde, followed by sucrose gradient dehydration (15%, 20%, 30%), with each gradient dehydration carried out for 12h at 4°C. The brain sample was then embedded by OCT, frozen in liquid nitrogen, sectioned at −25°C, 12um thickness. The slices were treated in 3% hydrogen peroxide for 15min, at room temperature, PBS washed three times and incubated by normal goat serum at room temperature, and then incubated with rabbit anti-mouse CD4 antibody and rabbit anti-mouse CD8 primary (Bioss, China; 1: 100, 4°C overnight) antibodies. Washed with PBS, stained with HRP conjugated secondary antibody, and incubated in DAB solution for color development. Four images were captured for each tissue section, the mean number of positive cells (number/mm2) was calculated by using IPP 6.0 software. The criteria of positive were the cytoplasm has brown particles.

### 2.8 Immunofluorescence

The tissue sections were fixed in 4% paraformaldehyde solution for 15 minutes and washed with distilled water for 5 minutes. Then blocked with 10% goat serum, 28°C for 1h. After blocking, the sections were incubated with primary antibody overnight at 4°C, washed 3 times with PBS buffer, 5 min each time. Then the sections were incubated with different fluorescently labeled secondary antibodies at 37°C for 1 h, washed with PBS buffer, and stained with DAPI solution for 15 min followed by PBS wash. The sections were then mounted with glycerol and covered with coverslips.

### 2.9 RT-PCR

Four hundred nanogram total RNA was employed for cDNA synthesis using ProtoScriptII First Strand cDNA Synthesis kit (TaKaRa,#RR036A) according to the manufacturer’s protocol. Forty ng cDNA was used as templates for RT-PCR using SYBR® Premix Ex Taq kit (TaKaRa, #RR820A) using LightCycler 96 (Roche, Switzerland). The sequences and efficiency of all primers were listed in (Supplementary Table 2), and HPRT/GAPDH was used as reference genes.

### 2.10 Western-Blot

Protein was extracted on ice in RIPA solution (Solarbio, #R0020), and quantified using a bicinchoninic acid protein assay kit (Solarbio, #PC0020). Forty μg protein was separated by 8% SDS/PAGE, transferred to nitrocellulose membranes (MILLIPORE, #HATF00010) at 120v for 90min. Membranes were blocked with 5% Skim Milk solution (Solarbio, #D8340) for 3h at 37°C, immunoblotted with primary antibody (listed in Supplementary Table 1) overnight at 4°C, washed, then blotted with a secondary antibody, washed, and imaged using ODYSSEY CLX (Clx-0519, LI-COR, Gene Company Limited, USA).

### 2.11 Mouse peripheral blood mononuclear cell collection

1. Anesthetize the mice with 3% chloral hydrate, take 700ul of whole blood from the mouse heart inject an anticoagulation tube containing heparin sodium, and mix well after adding the same amount of Hank’s solution.
2. Add 1/2 blood volume of the peripheral blood mononuclear cell separation solution (Solarbio #P6340) to the centrifuge tube, and suck mouse whole blood with a pipette, then slowly inject to the liquid surface of the stratification solution along the tube wall.
3. Put it in a centrifuge at 1500 r/min for 10 min. It can be seen that most of the albuginea-like mononuclear cells are suspended at the interface between the plasma and the stratified fluid.
4. Suck the white membrane cell layer with a straw, transfer it to a new centrifuge, add the same amount of Hank’s solution, mix the liquid, put it in a horizontal centrifuge at 1000r/min for 10min, discard the supernatant.
5. Repeat step 4 twice.
6. Add RPMI-1640 complete medium (1% penicillin-streptomycin solution + 10% FBS) to the pellet, mix well and transfer it to a 5 ml cell culture flask with complete medium, and cultivate in a CO2 incubator.

### 2.12 Statistical analysis

Statistical analysis was carried out by both the SPSS package and Prism6.0 software package. Tumor growth analysis was done by applying a linear mixed model using an online tool TumorGrowth(https://kroemerlab.shinyapps.io/TumGrowth/). Immune cell percentage in four groups was determined by ANOVA using Graphpad. Overall survival was defined as the time from glioma cell engraftment until the sacrifice of the animals. Survival curves were plotted using the Kaplan–Meier method and compared by log-rank test using SPSS. A p-value less than 0.05 was considered significant.

### 2.13 Institutional Review Board Statement

The study was conducted according to the guidelines of the Declaration of Helsinki and approved by the Institutional Review Board (or Ethics Committee) of Inner Mongolia Agricultural University.

## 3 Results

### 3.1 ITE synergized with PD1 antibody to suppress tumor growth and extend animal survival

To assess ITE’s anti-glioma effect, we established an orthotopic mouse glioma model by injecting GL261 cells intracranially, and monitoring tumor growth by in vivo imaging using a near-infrared fluorescent dye, IR783, which specifically targets tumor cells. Tumors became detectable 11 days after implantation, which was designated as day 1. (Figure1A-B). Animal body weights were monitored daily and treatment with ITE, PD1 antibody or the combination of ITE and PD1 antibody (designated as ITE+PD1) started on the first day of bodyweight drop(Figure1C). Animals were observed daily and imaged every three days, euthanized when the posterior tumor edge reaches the tragus of the ears for animal welfare. We fitted a linear mixed model using the online tool TumorGrowth to analyze the imaging data. The results showed that although the ITE+PD1 group has a trend of slower tumor growth compared with the DMSO control group(Figures 1E and 1F), the growth curve difference did not reach statistical significance(p>0.05). There were long-term survivors in the PD1 and ITE+PD1 groups, which were observed for 100~180days after euthanasia of all other animals, and found to be cured as the autopsy revealed(data not shown).

**Figure 1.** ITE synergized with PD1 antibody to suppress tumor growth and extend animal survival:(A) 1×10^5^ mouse glioma cells GL261 were injected into the brain of mice through a stereotaxic device to construct a tumor model; (B) Detect whether the model mouse is tumorigenesis: Left: H&E staining of mouse whole brain sections; Right: Inject the fluorescent dye IR783 (100nmol/mouse) into the intraperitoneal cavity of mice, and the fluorescent area is the enrichment of the tumor; (C) Four groups of drugs were given to the model mice: 10% DMSO/mouse, 100 mg/kg ITE/mouse, 80 μg PD1 antibody/mouse, 100 mg/kg ITE, and 80 μg PD1 antibody combined treatment group; (D) Post-treatment mice were intracranially injected with 100umol IR-783, Average tumor luminescence was determined using an IVIS spectrum imaging station, representative IVIS spectrum images of treated mice with intracranial GL261 at days 0, 3, 6 and 9 post-intracranial injection treatment; (E)Tumor volumes. (F)Trends of tumor growth extracted by applying a linear mixed model to the growth data.; (G) Survival analysis for wild-type C57BL/6 mice intracranially 1× 10^5^ GL261 cell s and receiving DMSO?> ITE PD1 and combination-therapy ITE with PD1, all treatment beginning at 14 days post-intracranial injection (post-ic.).

### 3.2 ITE synergized with PD1 antibody to increase cytotoxic immune cells infiltration in glioma

Flow cytometry analysis was carried out utilizing glioma and spleen tissue from different treatment groups. The gating and cell identification strategy are shown in Figure2A. Firstly, increased CD45+ cells percentage was found in the ITE+PD1 group in both glioma tissue and spleen, showing more immune cell infiltration. Specifically, the cytotoxic CD4-CD8+T cells percentage was significantly increased in the ITE+PD1 group (Figure2) compared to the DMSO and ITE groups, while the PD1 group was in-between the DMSO and ITE+PD1 groups, not reaching statistical significance from either group. Immunohistochemistry (IHC) and immunofluorescent (IHF) staining of glioma tissue confirmed such increase and further revealed that CD8+ T cells infiltrated deeper into the glioma tissue upon ITE treatment (Figure 2C). In contrast, another cytotoxic immune cell population, NK and NKT cells, marked by Nkp46, was not significantly changed in either glioma or spleen (Supp Figure 1).

**Figure 2.** ITE synergized with PD1 antibody to increase cytotoxic immune cells infiltration: (A-B) NK cell and adaptive cell (CD8, CD4, TH17, Treg) gating strategy; (c-g) The effection of immune cell in brain tumor and spleen with the treatment of DMSO, ITE, PD1 antibody and ITE +PD1 antibody; (C) CD45+ cells percent in brain tumor and spleen; (D) NK+NKT cell percent in brain tumor and spleen; (E) CD8+T percent in brain tumor and spleen; (F) Immunofluorescence and immunohistochemical staining of frozen slice from the brain tumor to detect T-cell antigens CD8; blue: cell nucleus, green: CD3, red: CD8; (G) CD4+T percent in brain tumor and spleen; (H) Immunofluorescence and immunohistochemical staining of frozen slice from the brain tumor to detect T-cell antigens CD4; (I-J) TH17 and Treg percent in brain tumor and spleen.

Increased CD8+T cells infiltration could be the result of increased peripheral CD8+T cells recruitment, or in situ expansion of T cells in the glioma tissue, so we examined the T cells in the spleen and found that CD4-CD8+T cells percentage was not significantly affected(Figer 2D), indicating that spleen was not the origin of these cytotoxic T cells. To assess the function of these tumor infiltrated CD8+T cells, the IFN-γ level was determined, which was significantly increased in the PD1 and ITE+PD1 groups (Figer 2E).

To identify the mechanism of the increased cytotoxic immune cell infiltration, we further analyzed major cell populations that regulate CD8+T cells infiltration. CD4+T cells were significantly increased in the ITE+PD1 group’s glioma tissue but not the spleen, indicating again a local expansion instead of the peripheral recruitment. We further analyzed the Th17/Treg ratio, a well-known regulator of tumor infiltrated CD8+T cells. Although showing a trend of increasing, Th17/Treg did not change significantly (Figure2G-H) in the brain, indicating that Th17/Treg was not the major factor boosting cytotoxic T cells infiltration.

### 3.3 ITE reduced MDSC infiltration in the brain tumor

MDSCs exert multiple immune suppressive effects and are further divided into Ly6ChighLy6G-(designated as M-MDSC) and Ly6ClowLy6G+ (designated as PMN-MDSC) cells in mice (Figure3A). It’s worth noting that another group of pro-tumor myeloid cells, neutrophils, are also Ly6ClowLy6G+. M-MDSC percentage was significantly reduced in the ITE+PD1 group (Figure3C), while PMN-MDSC+neutrophil cells percentage was significantly reduced in the ITE group (Figure 3C). Accordingly, the sum of M-MDSC, PMN-MDSC+ neutrophils (Designated as MDSC) percentage was significantly reduced in both ITE and ITE+PD1 groups. Immunofluorescence of the mouse glioma tissue with Gr-1 and CD11b antibodies further confirmed this observation (Figure3D). Moreover, the pattern of MDSC infiltration was changed, too: MDSCs distributed more at the peripheral of the glioma tissue compared to the DMSO or PD1 group, just in contrast to the CD8+T infiltration pattern. No significant changes were observed in the spleen (Figure3E), indicating that ITE probably inhibited the MDSC infiltration on steps other than recruitment from peripheral tissue.

**Figure 3.** ITE reduced MDSC infiltration in the brain tumor: (A) PMN-MDSC, M-MDSC, and MDSC cell gating strategy; (B) The linear relationship between the ratio of M-MDSC and the survival of model mice; (C) Proportion of M-MDSC, PMN-MDSC and MDSC cells in brain tumor tissue treated by DMSO, ITE, PD1 antibody and ITE combination with PD1 antibody; (D) Detection of antigens MDSC in brain tumor with the treatment of DMSO, ITE, PD1 antibody, ITE combination with PD1 antibody by Immunofluorescence; blue: cell nucleus; green: CD11B; red: Gr; Merge: MDSC (*P<0.05,**P<0.01); (E) Proportion of M-MDSC, PMN-MDSC and MDSC cells in spleen treated by DMSO, ITE, PD1 antibody and ITE combination with PD1 antibody.

As a preliminary functional analysis of MDSC cells, we plotted MDSC percentage against mouse survival days and found that the MDSC percentage negatively correlated with survival (r=0.63), further revealing the important roles played by MDSC (Figure 3B) in this model.

### 3.4 ITE treatment significantly downregulated IL11 in GL261 cells and glioma tissue

To reveal the possible mechanism of ITE’s regulation on MDSC, we firstly examined AHR protein level in GL261 cells and glioma tissues and found that AHR protein level was slightly increased upon ITE treatment of GL261 cells in vitro (Supp Figure Figure 4A), and remained unchanged in mouse glioma tissues of all groups (Figure 4B), ruling out the possibility of ITE’s effect depending on diminished AHR level.

**Figure 4.** ITE treatment significantly downregulated IL11 in GL261 cells: (A) Western-Blot to detect the protein level of AHR in GL261 cells treated with DMSO and ITE (0.1nM, 10nM, 10000nM) for 20 h (*P<0.05); (B) Western-Blot to detect the protein level of AHR in brain tumor treated with DMSO, ITE, PD1antibody and ITE combination with PD1 antibody; (C-D) Q-PCR and Western-Blot to detect the mRNA and protein level of IL6 and IL11 in GL261 cells treated with DMSO and ITE (0.1nM, 10nM, 10000nM) for 20 h (*P<0.05,**P<0.01); (E-F) Western-Blot and Immunofluorescence to detect the IL6 and IL11protein level in brain tumor treated with DMSO, ITE, PD1antibody and ITE combination with PD1 antibody; DAPI: cell nucleus, Green: IL6, Red:IL11 (*P<0.05,**P<0.01); (G-H) Western-Blot to detect the protein level of IL6 and IL11 in GL261 cells treated with Kyn (1μM, 10μM) and 1μM KYN combination with ITE (10nM, 100nM, 1000nM) for 20h (*P<0.05, **P<0.01); (I) PBMCs was collected from the blood of normal C57BL/6 mice and treated with IL-11 (10nM) and ITE (100nM) and 10 nM IL11 combination with 100 nM ITE for 5 days. The number and morphology of PBMC were recorded under a microscope; (J) Flow cytometry to detect the proportion of M-MDSC in PBMC treated by DMSO, 10nM IL-11, 100nM ITE, and 10 nM IL11combination with100 nM ITE.

RNA seq data was collected from ITE or DMSO treated U87MG cells and GL261 cells (data available at GEO database U87MG reference PRJNA788010, GL261 reference PRJNA789328). ITE’s effects on gene expression profiles of interleukins, chemokines, and other immune-regulatory factors were checked in the RNA-seq data(summarized in supplemental Table3). We searched for factors that are known to regulate MDSCs with concordant changes and found that ITE reduced IL6 family members, IL6 and IL11 mRNA levels in GL261 cells (Figure4C-D). Higher ITE doses significantly reduced IL11 protein levels in cultured GL261 cells (Figure4D). In contrast, the known AHR target, IL6 protein level remained unchanged, which was inconsistent with the mRNA level, indicating additional regulation in the post-transcription level of IL6.

To confirm that IL11 was an AHR target, cells were subjected to KYN (1μM, 10μM), and both IL6 and IL11 increased significantly (Figure4G). We further tested ITE’s effects in presence of KYN and found that with 1uM KYN, IL6 and IL11 expression was inhibited by ITE (Figure 4H), suggesting that ITE might inhibit IL11 in tumor tissue in presence of endogenous KYN. Accordingly, in mouse glioma tissue, IL11 level was inhibited in both the ITE and the ITE+PD1 group, with the ITE+PD1 group reaching statistical significance (Figure 4E-F).

IL11 can induce PBMC differentiation into MDSC, and we tested whether ITE can block this process in vitro. As expected, the addition of ITE to the cultured PBMC with IL11 significantly reduced M-MDSC percentage, revealing ITE’s ability to block IL11 induced PBMC differentiation toward MDSC (Figure 4I-J). ITE’s inhibition effect reaches statistical significance only with the addition of IL11, indicating that elevated IL11 is a prerequisite for ITE’s inhibition of MDSC differentiation.

In glioblastoma, tumor-associated microglia and macrophages secreted IL11 to activate the STAT3-MYC pathway in glioma cells (PMID: 33846242), so we tested ITE’s effects on STAT3 in glioma cells. Phosphorylated stat3(pStat3) was significantly inhibited in ITE-treated GL261 cells (Figure 5A), confirming ITE’s blocking effects of the IL11-STAT3 axis in cultured glioma cells. In contrast, 10μM, 100μM Kyurenine significantly increased phosphorylation of STAT3 (Figure 5B), in a dosedependent manner. With the presence of 1μM KYN, 100nM ITE significantly inhibited STAT3 (Figure 5C), which is following ITE’s IL11 inhibiting effects in presence of 1uM KYN. STAT3 is a critical mediator of immune suppression, so we further explored how ITE affected STAT3 in the glioma tissue. Unexpectedly, the pStat3 level increased in the ITE+PD1 group (Figure 5D), probably due to other factors upstream of STAT3.

**Figure 5.** ITE treatment significantly downregulated pSTAT3 in GL261 cells : (a) Western-Blot to detect the protein level of STAT3 and pSTAT3 in GL261 cells treated with DMSO and ITE (10nM, 100nM, 1000nM) for 20h (*P<0.05, **P<0.01); (b-c) Western-Blot to detect the protein level of STAT3 and P-STAT3 in GL261 cells treated with Kyn (1μM, 10μM) and 1μM KYN combination with ITE (1nM, 10nM, 100nM) for 20h (*P<0.05, **P<0.01); (d) Western-Blot to detect the protein level of STAT3 and P-STAT3 in brain tumor treated with DMSO, ITE, PD1, ITE+PD1 combination.

## 4 Discussion

Recently, several AHR antagonists have been studied as drug candidates. CH223191 has been reported to restrict a Treg-macrophage suppressive axis^[20]^. AHR inhibitor, BAY 2416964 inhibited tumor growth in mouse models and entered clinical trials recently^[21]^ both in the U.S and China. We reported that ITE, an endogenous AHR ligand usually classified as an agonist, can activate anti-tumor immunity when combined with PD1 antibodies, probably via suppressing the IL11-MDSC pathway. Previously ITE was reported to suppress immunity in experimental autoimmune encephalomyelitis (EAE) and experimental colitis^[21–23]^, and the discovery of ITE’s immune-activating effects via inhibiting MDSC in glioma highlighted the complexity of AHR signaling. Moreover, the discovery of IL11 as an AHR target provided new insights on how AHR might affect MDSCs.

The immune profile of the orthotopic GL261 mouse model by multi-color flow cytometry revealed that in the ITE+PD1 group cytotoxic CD4-CD8+T cells infiltrated more and deeper into glioma tissue^[24, 25]^, indicating the tumor microenvironment became “hot” with the treatment. While 80μg PD1 antibodies alone failed to show such an effect. Studies have shown that the CD8+ T cell infiltration in GBM is positively correlated with the long-term survival of GBM patients^[26]^. Accordingly, we observed that more mice were cured by the combination therapy compared to the PD1 group. The increased IFN-gamma also confirmed activated anti-tumor immunity.

One factor that may contribute to the opposite immune-modulatory effects of ITE in EAE and glioma could be AHR’s interaction partner having opposite effects. One such partner is the estrogen receptors. The fact that the prevalence of astrocytoma, a glioma subtype, is higher in men^[27]^, while the prevalence of multiple sclerosis (which is modeled by EAE) is higher in women^[28–30]^ suggested that estrogen played opposite roles in these two diseases. Estrogen receptors directly interact with AHR and modulate AHR’s function as a transcription factor. Other factors including HIF1alpha, RelA, RelB, E2F1, Rb, AR also interact with AHR and could modulate its effects^[31]^.

Another possible reason for ITE’s opposite effects in auto-immunity and the GL261 glioma model might be a different dosage regime. ITE suppressed immunity via inducing tolerogenic DCs and Tregs, at 200ug/mouse(equals to 10mg/kg for typical 20gram mice) in EAE, and the same dose also induced Tregs (10mg/kg, three or four days apart) in experimental colitis^[32]^. A higher dosage (100mg/kg) was applied at 48hr intervals in our study. Another high-affinity AHR ligand, FICZ, also has dose-dependent immune-modulating effects: 50ug/kg FICZ promoted, while 10mg/kg FICZ inhibited Th17 cells in an acute alloresponse model^[33]^. Due to the different and sometimes opposite effects of the same AHR ligand, AHR ligands have been proposed as selective modulators instead of agonist or antognist^[34]^.

For understanding the different effects of AHR ligands, Ehrlich et al demonstrated that the immune regulation effects of AHR ligands depend on dose and duration of AHR activation, gauged by activation of CYP1A1 transcription. In the acute alloresponse model, ITE’s induction of the CYP1A1 was transient, with a single dose of 10mg/kg or 40mg/kg ITE’s effect peaked at 4hr and diminished at 20hr. Two doses of 80mg/kg ITE administered 12hr apart or 40mg/kg administered every 6 hr could induce CYP1A1 to a comparable level of one dose of 15ug/kg TCDD, gauged by both activation of CYP1A1 transcription and inhibition of splenic CD8+T cells. It is possible that with our dose regime each dose can induce a high but transient transcriptional activity peak, which activates immunity in the glioma model, while the sustained activation with 80mg/kg every 12hr inhibited immunity in the alloreponse model, and EAE and experimental colitis model.

Accordingly, using an ultrasensitive reporter, Hoffman et al classified AHR ligands into two classes: potent AHR activators with sustained transcriptional activation and toxic responses, and mild AHR modulators with transient transcriptional conditioning and subtle therapeutic effects; ITE fits better as a mild AHR modulator^[35]^. Our RNA-seq data collected 24hr after adding ITE also supported ITE as an AHR modulator: CYP1A1 or other cytochrome p450 gene expression changes were not detected.

For immune-landscape changes, we also monitored tumor infiltrated CD4+T cells, especially the Th17 and Treg populations, for AHR is a master regulator for CD4+T cells development. For example, TCDD directly modulates both CD4+ and CD8+ T cells^[36]^, and KYN suppressed proliferation of T cells^[37]^ and memory CD8+T cells^[38].^ Moreover, in response to the PD1 antibody-induced IFN gamma increase, the IDO-KYN pathway inhibited CD8+T cells by controlling Treg differentiation to lower the Th17/Treg^[39]^. However, ITE+PD1 did not significantly change the Th17/Treg, so other factors regulating CD8+T cells were searched, which turned out to be down-regulation of the MDSC population in glioma tissue. ITE+PD1 significantly reduced the total MDSC and the M-MDSCs in the glioma tissue, while neither ITE nor PD1 alone reached statistical significance. MDSC but not Tregs was elevated in peripheral blood of GBM patients, and higher MDSC in recurrent tumors compared to primary tumors predicted poor prognosis in glioma^[40]^. MDSC is reported to mediate PD1 resistance via inhibiting CD8+ T and NK cells, therefore ITE’s MDSC inhibition effect may play a critical role in overcoming PD1 resistance. It’s worth noting that MDSC in glioma drives tumor growth in a sex-specific manner: depletion of PMN-MDSC only extended survival in female mice, while M-MDSC was predominant in males^[41]^. As a common practice, we have used all-female mice in our study^[42, 43]^, and observed that ITE significantly reduced PMN-MDSCs, a prognostic marker in female patients’ tumors. More studies in male mice are needed to determine ITE’s effects in males.

In response to sustained myeloid growth factors and inflammatory signals, MDSCs emerge in two steps: first, they expand and condition in bone marrow and spleen, then convert/differentiate from neutrophils or monocytes in cancer tissue. ITE could regulate one or more steps in this process. It’s known that TCDD leads to massive mobilization of MDSC with immunosuppressive activity through the regulation of CXCR2 and miR-150-5p and miR-543-3p^[44]^. ITE could also regulate MDSCs’ functions, as AHR expressing oral squamous carcinoma cells induced expression of checkpoint molecules PDL1 and CD39 in MDSC^[45]^.

Although we did monitor M-MDSC and PMN-MDSC in the spleen, our data is not sufficient for discerning whether ITE affected the expansion, mobilization, or conversion/differentiation of MDSCs, as we lacked the data on neutrophils and monocytes. Spleen Ly6C+Ly6G-cells could be either M-MDSC or the newly identified PMN-MDSC progenitors, monocyte-like progenitors of granulocyte (MLPG)^[46]^. Further analysis with additional markers is required to clarify how ITE affected each MDSC species.

Since cancer cells are the source of signals attracting myeloid cells, we sought to identify factors that were both ITE-AHR targets and regulate MDSC and discovered IL11. Among multiple factors that up-regulate MDSC, such as target genes of STATs family members (STAT1, STAT3, STAT5, STAT6), M-CSF, G-CSF, VEGF, IL-1, IL-4, IL-6, IL-13, and IFN gamma^[17]^, several are also AHR targets, but they did not show significant changes in RNA-seq data of ITE treated glioma cells. So we started from cytokines /factors that were changed upon ITE treatment and discovered IL6 and IL11, with the latter further confirmed at the protein level. Although IL-6 is a well-known AHR target, its family member IL11 has not been associated with AHR before, and the discovery of ITE and KYN’s opposite regulation of IL11 supported it as an AHR target in glioma.

IL11 activates the STAT3-MYC signaling pathway in GBM cells, and accordingly, we observed reduced STAT3 phosphorylation upon ITE treatment in cultured glioma cells. The overall STAT3 phosphorylation level was increased in the ITE+PD1 group of glioma tissue, though. The following factors might contribute to this: firstly, glioma cells responded to other upstream factors of STAT3 that existed in tissue but not in the cell culture medium; secondly, non-cancerous cells such as macrophages responded to ITE by activating STAT3. Cell-specific knock-down experiments can further clarify the roles of STAT3 in MDSC infiltration. Activated STAT3 in the ITE+PD1 group suggested that STAT3 inhibitor shall be explored as an addition to the current combination reported here, specifically in those gliomas with a STAT3 gene signature ^[47]^ Finally, due to intrinsic differences in the mice and human AHR pathway, extrapolation of research results obtained in mouse gliomas to GBM patients requires caution.

## Supporting information

Table 1 Primer list

Table 2 Primary antibody list

table 3 RNAseq data of immunelate genes

## 6 Author Contributions

Conceptualization, Fang Wan, methodology, Lijiao Zhao, Fang Wan, software, Qiuting Shu, Yunlong Ma, validation, Lijiao Zhao, Yunlong Ma; formal analysis, Hui Sun, Lijiao Zhao, investigation, Jing Lu; resources, Pei Gong Fanhua Meng.; data curation, Yunlong Ma; writing—original draft preparation, Lijiao Zhao.; writing—review and editing, Fang Wan; visualization, Yunlong Ma; supervision, Fang Wan; project administration, Fang Wan; funding acquisition, Fang Wan. All authors have read and agreed to the published version of the manuscript.

## 8 Acknowledgments

Thanks to Jun Dong M.D. of Soochow University for his free teaching us the technology of in situ glioma modeling in mouse brain. Thanks to the National Natural Science Foundation of China for financial support for this research. Thanks to the Inner Mongolia Natural Science Foundation Committee for financial support for this research.

---

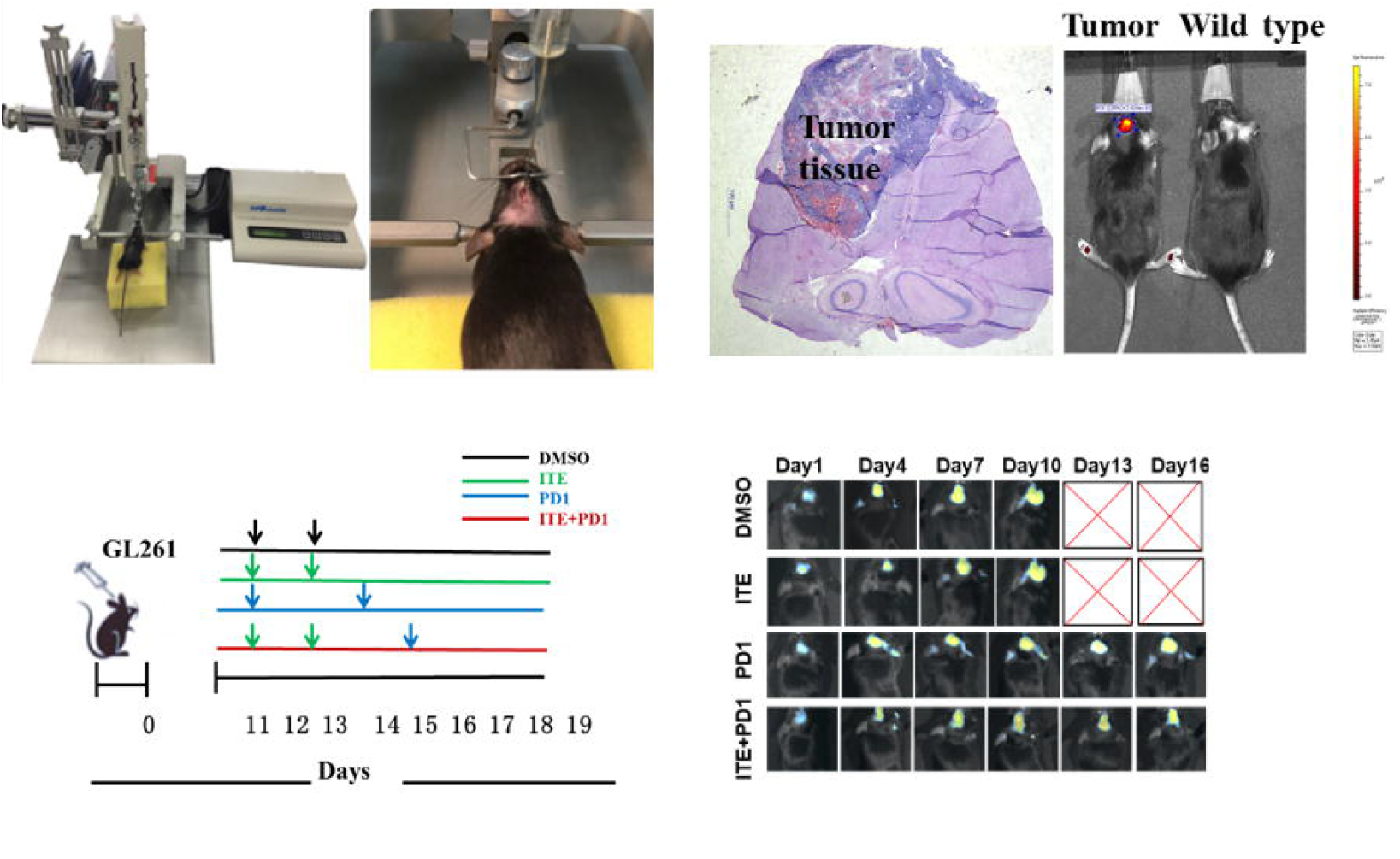
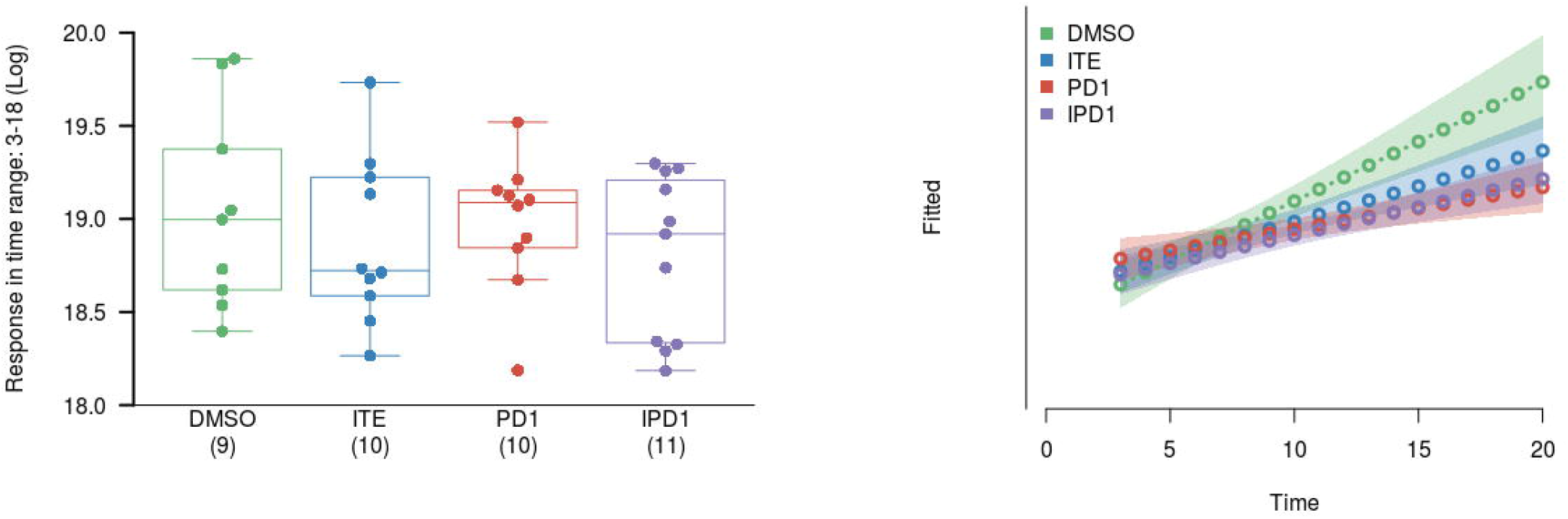
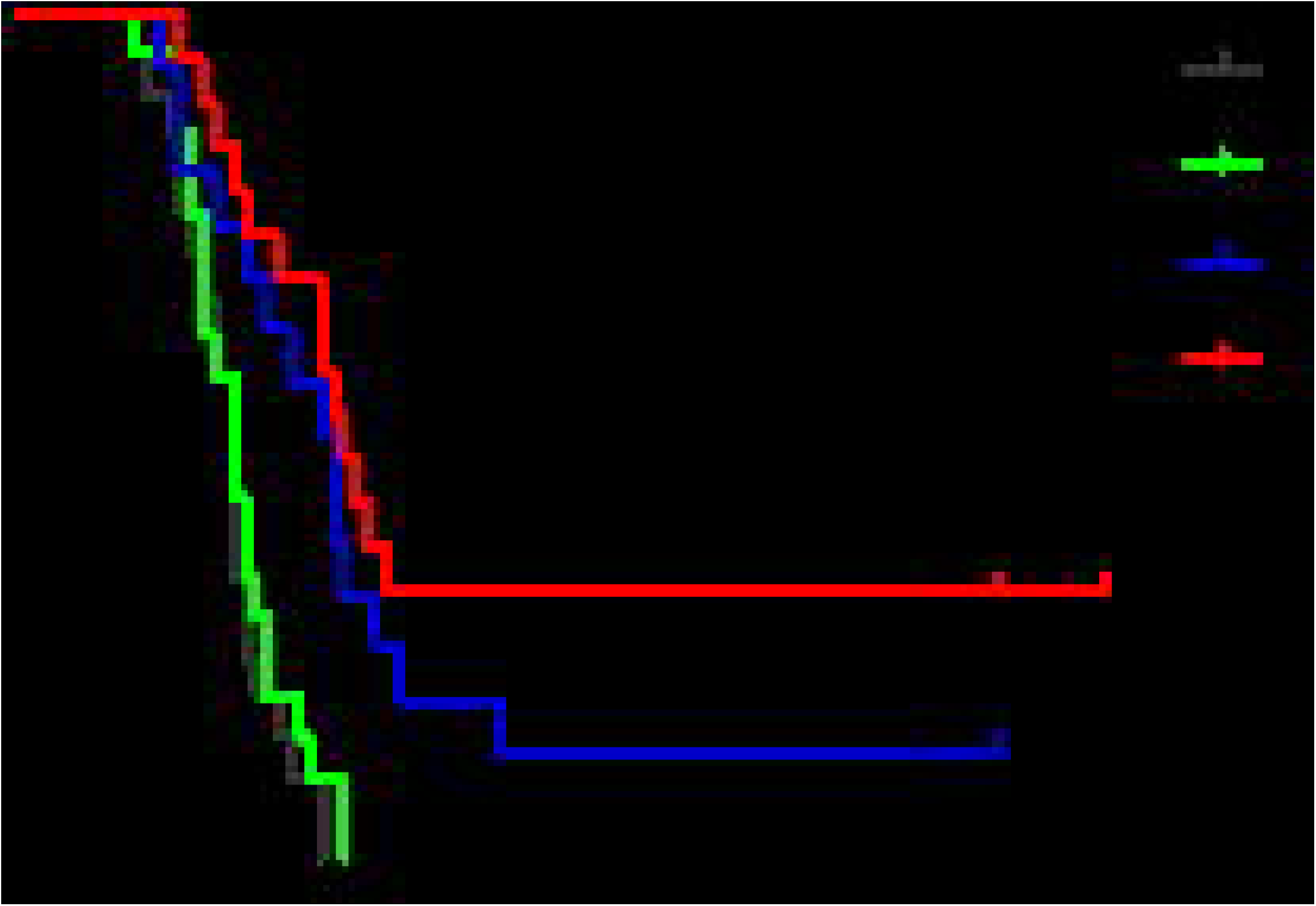
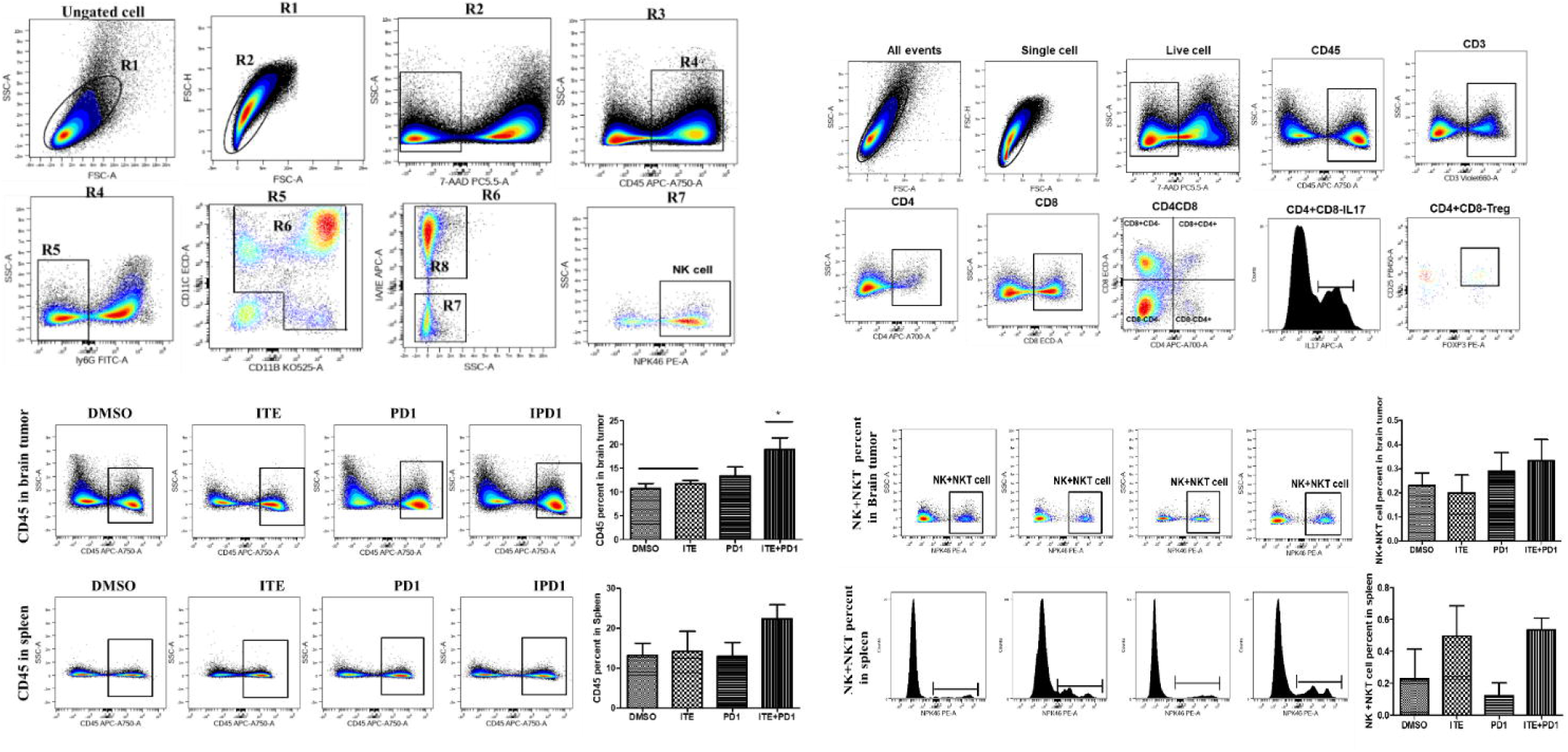
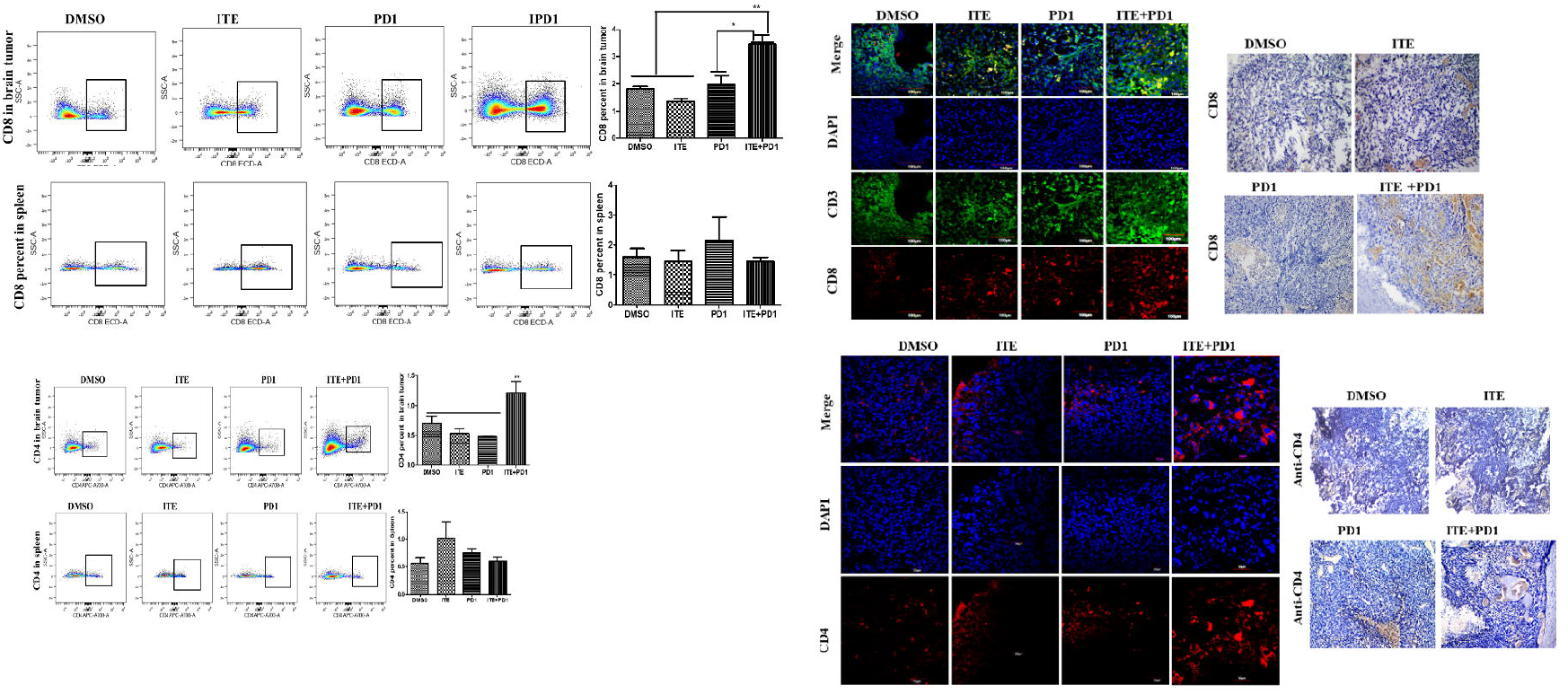
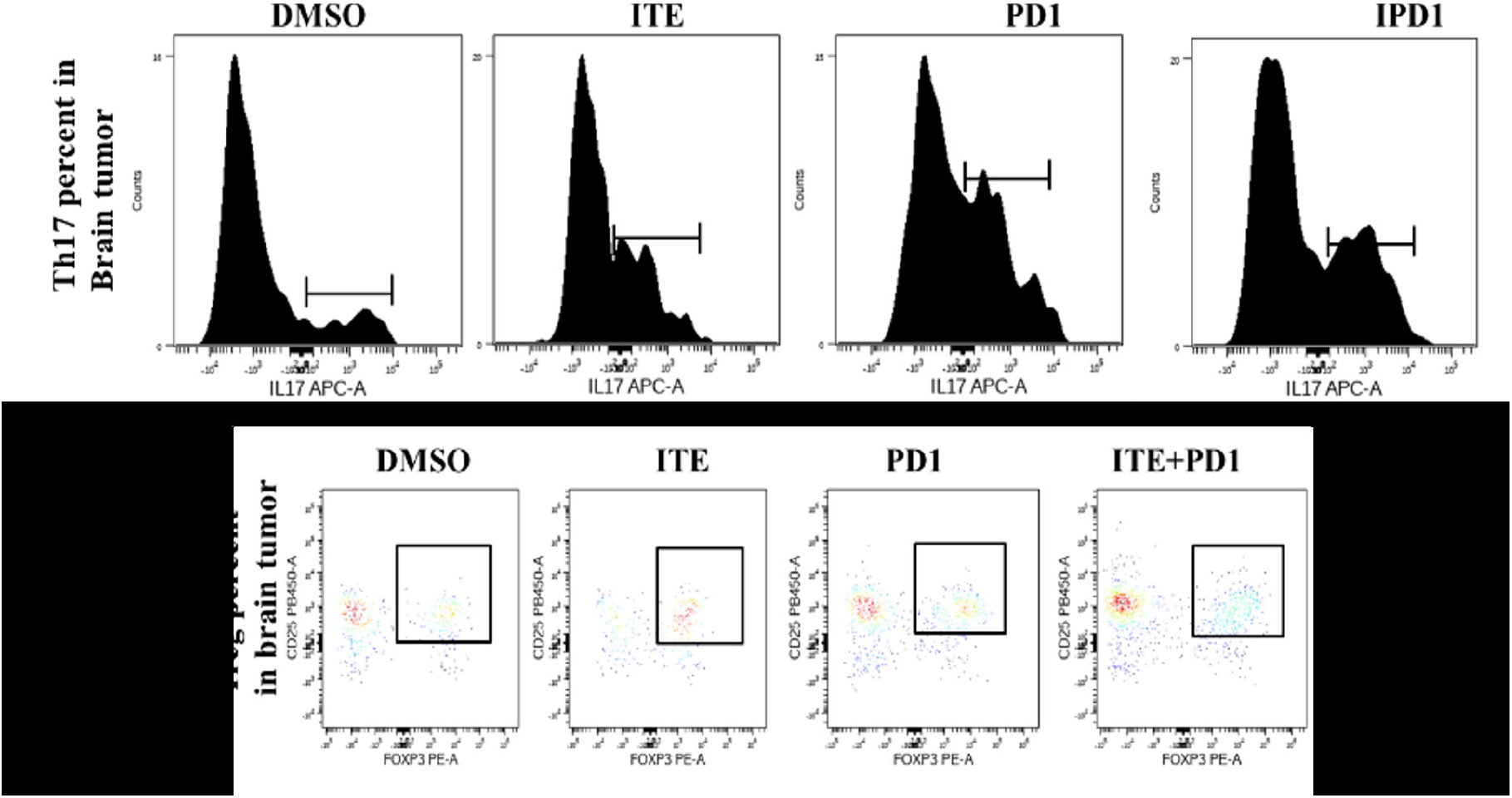
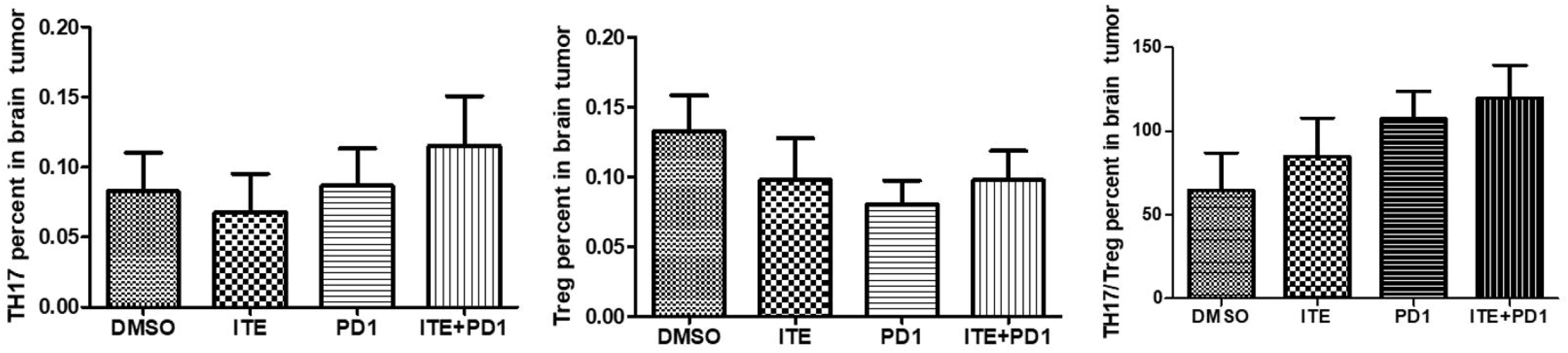
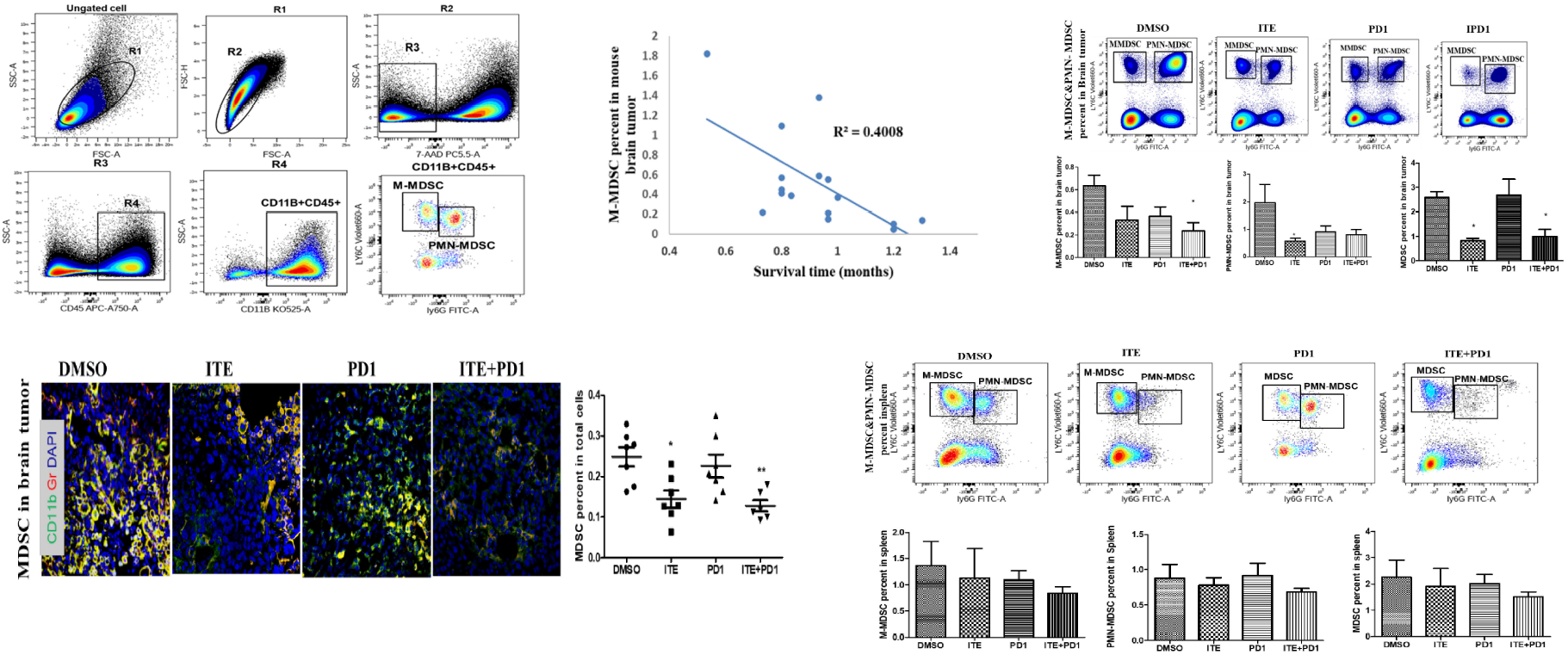
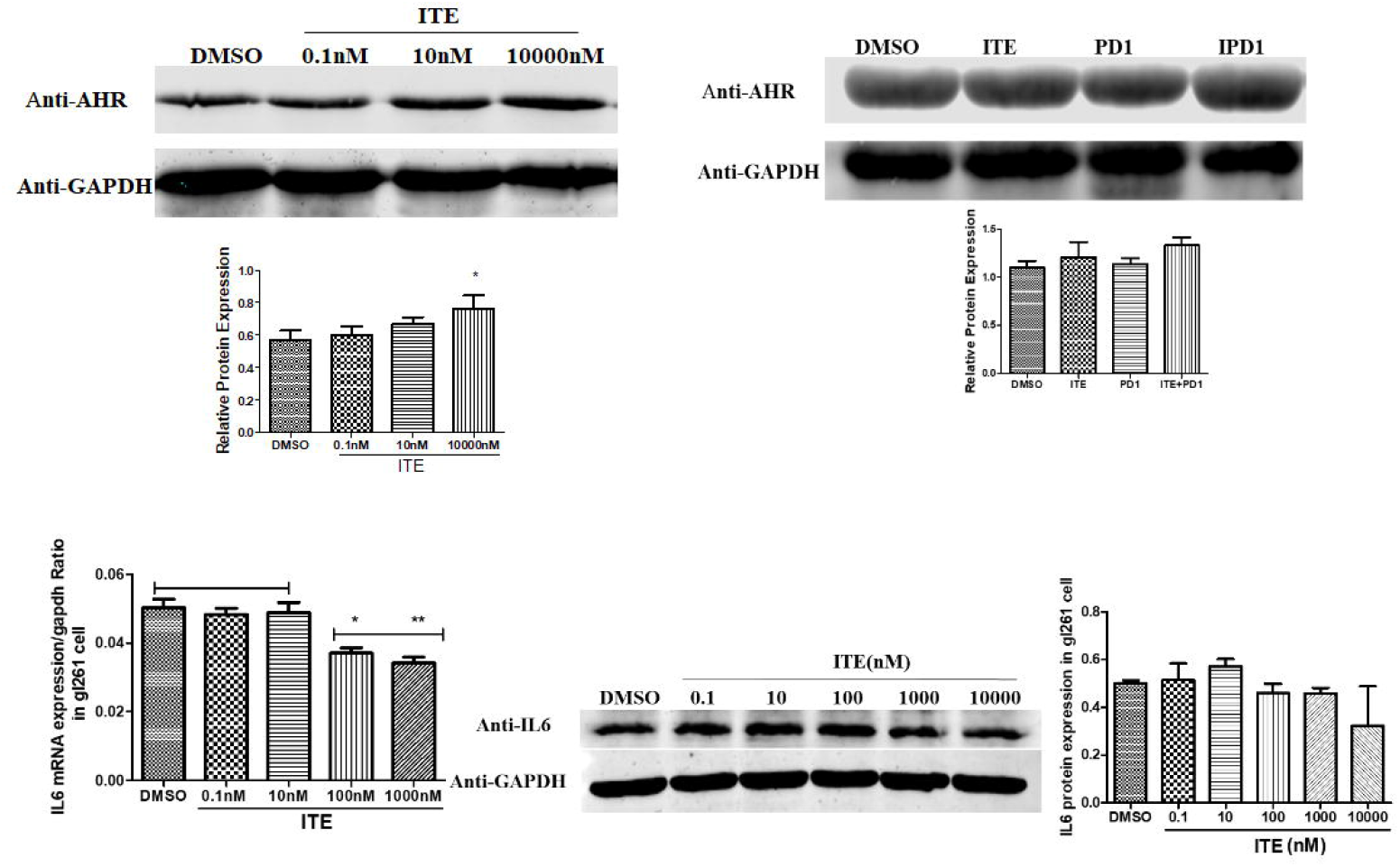
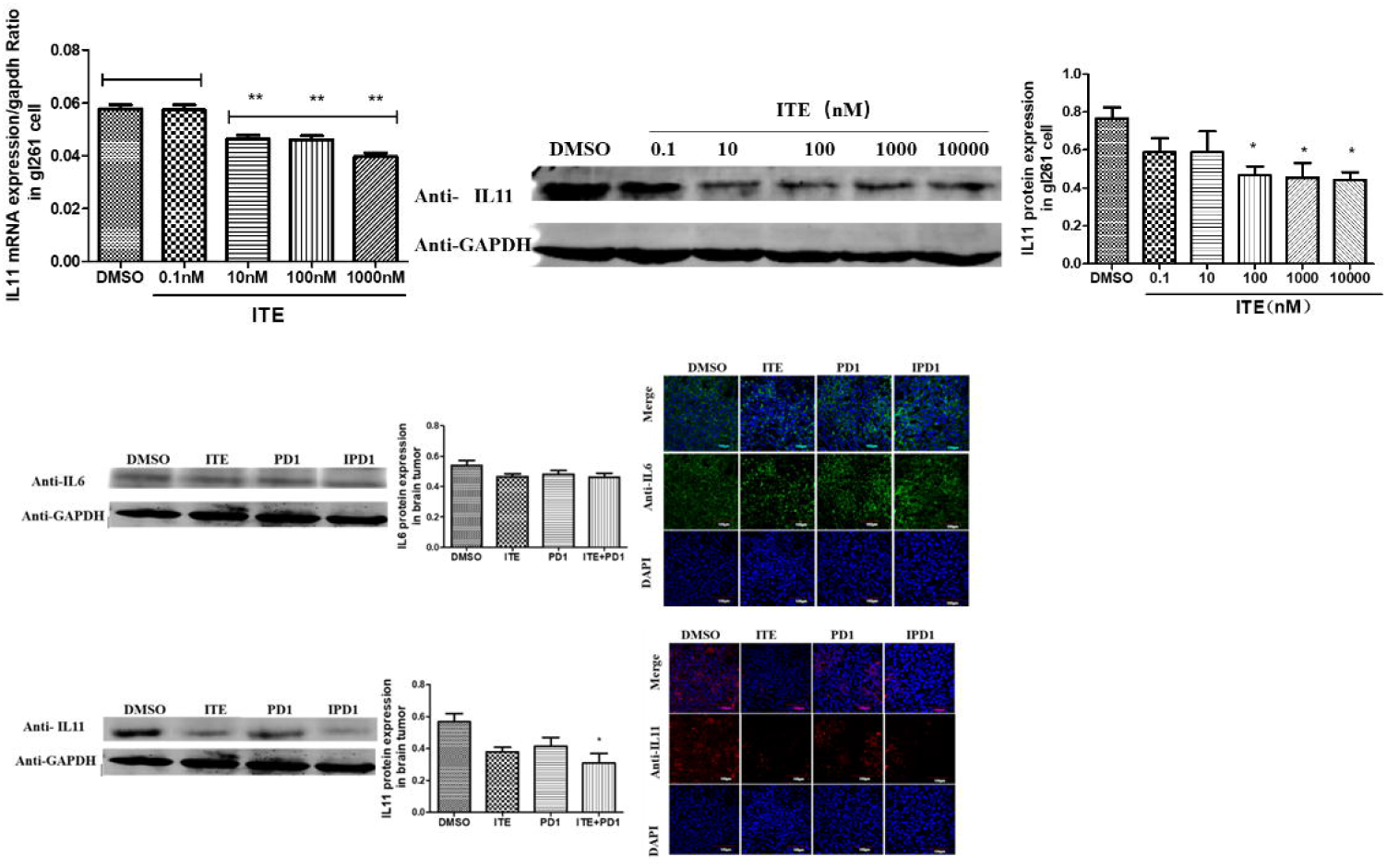
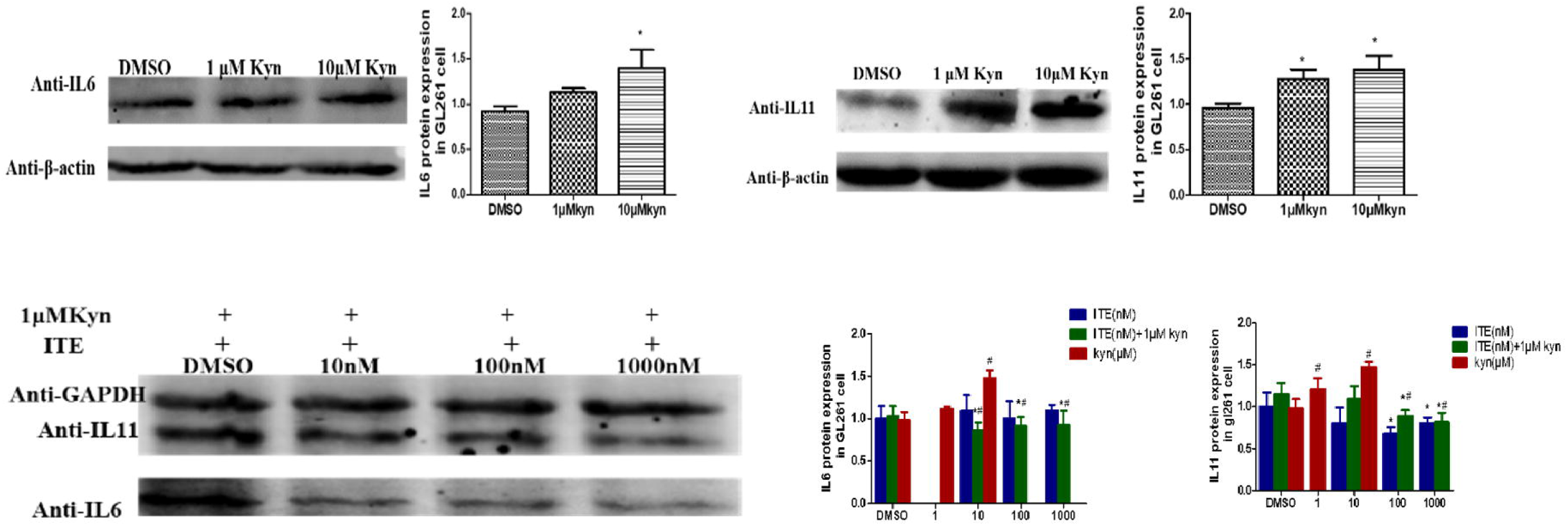
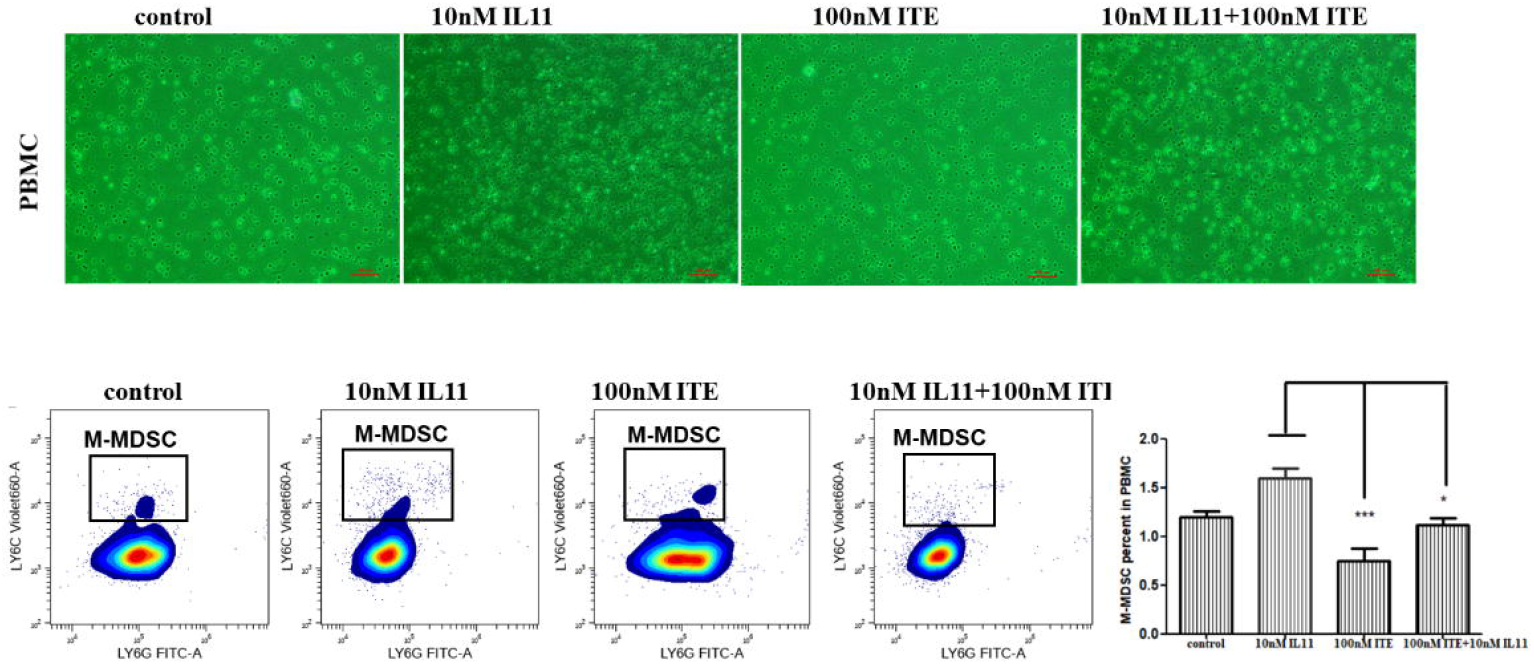
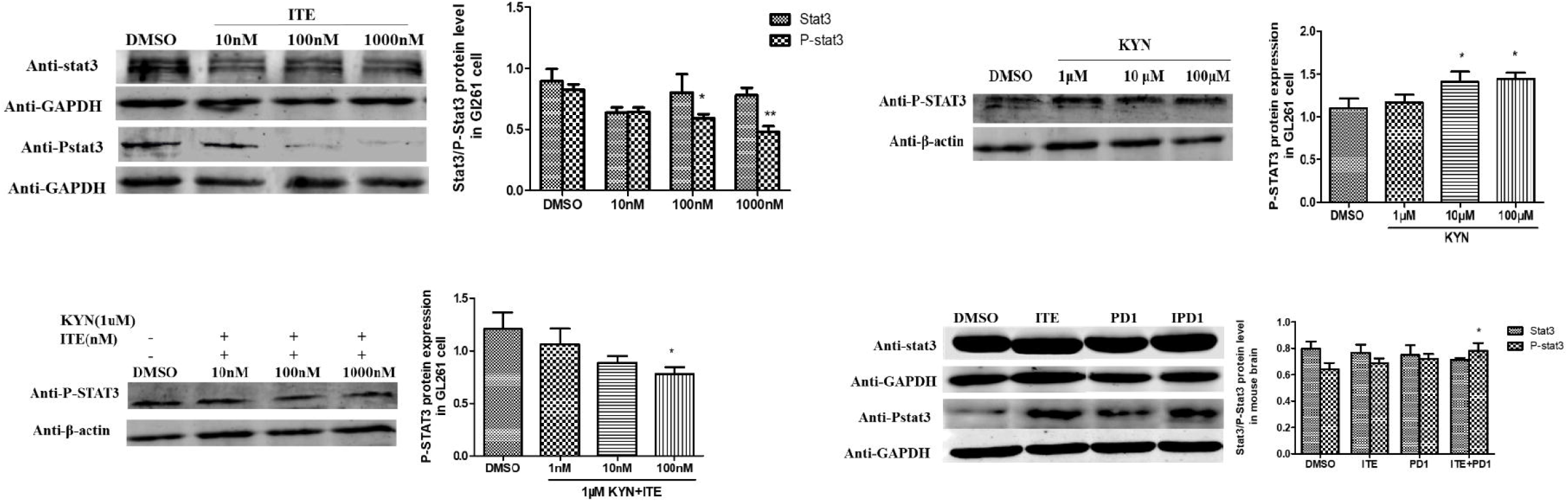

