## Supplementary material for "AHR agonist ITE boosted PD1 antibody’s effects by inhibiting myeloid-derived cells suppressive cells in an orthotopic mouse glioma model": Table 1 Primer list

Table.2 Primer list

| Gene | Forward primer | Reverse primer |
| --- | --- | --- |
| PTK2(human) | ACATTATTGGCCACTGTGGATGAG | GGCCAGTTTCATCTTGTTGATGAG |
| HPRT (human) | TGACACTGGCAAAACAATGCA | GGTCCTTTTCACCAGCAAGCT |
| MYH9 (human) | ATCTCGTGCTATCCGCCAAG | GTTGTACGGCTCCAACAGGA |
| Myh9(mouse) | ACGCCAAGACGGTGAAGAAT | CTTGGCGGATAGCACGAGAT |
| COCL1(human) | TCTGCGACAACGGCAAGGTG | GACGCCGGTGGTTTCTTGGT |
| ITGA5(human) | GCCTGTGGAGTACAAGTCCTT | AATTCGGGTGAAGTTATCTGTGG |
| ITGB5(human) | CAGGTGGAGGACTATCCTGTG | GTGCCGTGTAGGAGAAAGGAG |
| Vav3(mouse) | TTACACGAAGATGAGTGCAAATG | CAACACTGGATAGGACTTTATTCATC |
| VAV3(human) | ACGGACCAATGGACTGCG | TTCTGCCCTGCCAAAACA |
| Gapdh(mouse) | AACTTTGGCATTGTGGAAGG | GGATGCAGGGATGATGTTCT |
| GAPDH(human) | TCATTGAGCCCTTCCACAATG | GGTGTGAACCACGAGAAATATGAC |
