## Supplementary material for "AHR agonist ITE boosted PD1 antibody’s effects by inhibiting myeloid-derived cells suppressive cells in an orthotopic mouse glioma model": Table 2 Primary antibody list

| Antibody | Brand |
| --- | --- |
| CD8 | Abcam |
| CD4 | Abcam |
| STAT3 | Abclonal |
| STAT3 phosphorylation | SAB |
| LY6G | CST |
| Gr | CST |
| IL6 | Abclonal |
| IL11 | Abclonal |
| β-actin | Protein-tech |
| GAPDH | Protein-tech |
| a-tubμlin | Protein-tech |
| Goat anti-mouse fluorescent secondary antibody-488 | Abcam |
| Goat anti-mouse fluorescent secondary antibody-633 | Abcam |
| Goat anti- rabbit fluorescent secondary antibody | Abcam |
| Goat anti-mouse fluorescent secondary antibody-800W | Li-COR Bioscience |
| Goat anti- rabbit fluorescent secondary antibody-800W | Li-COR Bioscience |
| 7-AAD | Bio-Legend |
| CD45 | Bio-Legend |
| CD3 | Bio-Legend |
| CD4 | Bio-Legend |
| CD8 | Bio-Legend |
| CD25 | Bio-Legend |
| IL17A | BD |
| FOXP3 | BD |
| LY6G | Bio-Legend |
| CD11C | Bio-Legend |
| IA/IE | Bio-Legend |
| LY6C | Bio-Legend |
| NPK46 | Bio-Legend |
| CD11B | Bio-Legend |
| CD24 | Bio-Legend |
| CD64 | Bio-Legend |

Table 1 Primary antibody list
